# A large-scale circuit mechanism for hierarchical dynamical processing in the primate cortex

**DOI:** 10.1101/017137

**Authors:** Rishidev Chaudhuri, Kenneth Knoblauch, Marie-Alice Gariel, Henry Kennedy, Xiao-Jing Wang

## Abstract

We developed a large-scale dynamical model of the macaque neocortex based on recent quantitative connectivity data. A hierarchy of timescales naturally emerges from this system: sensory areas show brief, transient responses to input (appropriate for sensory processing), whereas association areas integrate inputs over time and exhibit persistent activity (suitable for decision-making and working memory). The model displays multiple temporal hierarchies, as evidenced by contrasting responses to visual and somatosensory stimulation. Moreover, slower prefrontal and temporal areas have a disproportionate impact on global brain dynamics. These findings establish for the first time a circuit mechanism for “temporal receptive windows” that are progressively enlarged along the cortical hierarchy, extend the concept of working memory from local to large circuits, and suggest a re-interpretation of functional connectivity measures.

**Conflict of Interest:** The authors declare no competing financial interests.

**Acknowledgements:** We thank Nikola Markov and John Murray for discussions. This work was supported by ONR grant N00014-13-1-0297 and NIH grant R01MH062349 (X.-J.W.) and by ANR-11-BSV4-501 and LABEX CORTEX (ANR-11-LABX-0042) of Université de Lyon, program “Investissements d’Avenir” (ANR-11-IDEX-0007) operated by the French National Research Agency (H.K.). The connectivity data is available at www.core-nets.org.

## Introduction

The receptive field is a central concept in Neuroscience, defined as the spatial region over which an adequate stimulus solicits rigorous response of a neuron (Sherrington, 1906). In the primate visual cortical system, starting with V1, the receptive field size of single neurons progressively enlarges along a hierarchy (Hubel and Wiesel, 1962; Hubel, 1988; Wallisch and Movshon, 2008). As a result, higher areas can integrate stimuli over a greater spatial extent, which is essential for such functions as size-invariance of object recognition in the ventral (“what”) stream for visual perception (Kobatake and Tanaka, 1994).

Accumulating evidence suggests that the brain also displays a hierarchy in the temporal domain. This allows neurons in higher areas to respond to stimuli spread over a greater temporal extent and to integrate information over time, while neurons in early sensory areas rapidly track changing stimuli. In human studies, preserving the short timescale structure of stimuli while scrambling the long timescale pattern changes responses in association areas much more than in early sensory areas (Hasson et al., 2008; Lerner et al., 2011; Honey et al., 2012; Gauthier et al., 2012). Notably, Honey et al. (2012) found that cortical areas sensitive to long time structure in the stimulus also show slower decays in their temporal autocorrelation (and hence slower dynamics). In the macaque, Murray et al. (2014) found a hierarchical organization in the timescales of spontaneous fluctuations of single neurons across 7 cortical areas. Moreover, an area’s position in the temporal hierarchy was well-predicted by its position in an anatomical hierarchy derived from long-range projection patterns (Felleman and Van Essen, 1991). Similarly, temporal correlations in neural activity reveal slower decay rates in the frontal eye fields compared to area V4 (Ogawa and Komatsu, 2010), the timescales of reward memory lengthen when moving from parietal to dorsolateral prefrontal to anterior cingulate cortex (Bernacchia et al., 2011), and, more generally, persistent activity after a brief stimulus has been shown to last for seconds, even across inter-trial intervals, in association areas including prefrontal cortex (Amit et al., 1997; Histed et al., 2009; Curtis and Lee, 2010). Finally, given that the brain needs to represent the environment on multiple timescales, normative theories of predictive coding suggest that a hierarchy of timescales would allow animals to form a nested sequence of predictions about the world (Kiebel et al., 2008).

What underlying neurobiological mechanisms might give rise to such a range of temporal dynamics? In particular, spatial patterns of convergence can produce increasing receptive field sizes in the visual hierarchy. Are there basic anatomical motifs that produce a hierarchy of timescales?

Here we report a large-scale circuit mechanism for the generation of a hierarchy of temporal receptive windows in the primate cortex. This hierarchy naturally emerges in a dynamical model of the macaque cortex that we developed based on a recently published quantitative anatomical dataset containing a directed and weighted connectivity matrix for the macaque neocortex (Markov et al., 2011; 2013b; 2014a; Ercsey-Ravasz et al., 2013). These data were obtained using the same experimental conditions and measures of connectivity, thereby ensuring a consistent database (Kennedy et al., 2013), and contain information both on the number of projections between areas and on their laminar origins. Based on a separate anatomical study (Elston, 2000; Elston et al., 2011), we introduced heterogeneity across cortical areas in the form of a gradient of excitatory connection strengths. Strong recurrent excitation has been proposed as a mechanism by which prefrontal cortex could implement “cognitive-type” computations, like information integration and memory-related delay activity; we hypothesized that differences in recurrent excitation might play an important role in the generation of a temporal hierarchy.

The model thus incorporates anatomically-constrained variation in both within-area and inter-areal connectivity, and allows us to probe the interplay of local microcircuitry and long-range connectivity that underlies a hierarchy of timescales. Using different sensory inputs we demonstrate the existence, in our model, of multiple dynamical hierarchies subserved by a single integrated global and local circuit. We then investigate the implications of local circuit heterogeneity for macroscopic dynamics measured by functional connectivity (i.e. correlations in activity across areas). Here we find a disproportionate role for slow dynamics in the prefrontal and other association cortices in shaping resting-state functional connectivity. This role is not predicted simply by long-range connections, suggesting that interpretations of brain imaging data will need to be revised to account for inter-areal heterogeneity.

While we have used the model to investigate the origin of a hierarchy of timescales (and implications for measures such as functional connectivity), this architecture can serve as a platform for future large-scale dynamical models relating anatomical connectivity to dynamical specificity and the functional roles of cortical areas. In particular, most previous statistical analyses of cortical connectivity (Bullmore and Sporns, 2009; Sporns, 2014) and computational models (Ghosh et al., 2008; Deco and Corbetta, 2011; Honey et al., 2007; 2009; Deco et al., 2014) have lacked comprehensive high-resolution data, relying either on collating qualitative tract-tracing data across disparate experiments and conditions or on diffusion tensor imaging (DTI), which is noisy and cannot reveal the direction of a pathway. Moreover, such models typically treat cortical areas as identical nodes in a network, distinguished only by their input and output connection patterns but not by local properties or computational capabilities. While this approach is reasonable for certain purposes, it is doubtful that functional specialization of cortical areas can be elucidated without taking heterogeneity into consideration. Our model provides a framework within which to explore how dynamical and functional specialization can emerge from inter-areal pathways coupled with local circuit differences.

## Results

We developed a large-scale model of the primate cortex in three steps. First, we used a recent connectivity dataset for the macaque neocortex (Markov et al., 2014a), specifically designed to overcome the limitations of collated anatomical data sets. In contrast to previously existing datasets, the data we use were collected by the same group under similar experimental conditions, with quantitative measures of connectivity that were designed to be comparable across experiments. As summarized in Figure 1, the connectivity weights are directionally specific and quantify connections between 29 widely-distributed cortical areas, with 536 connections observed to exist and inter-areal connection strengths spanning five orders of magnitude. The presence or absence of all projections in this network has been established; thus there are no unknown pathways.

**Figure 1.**
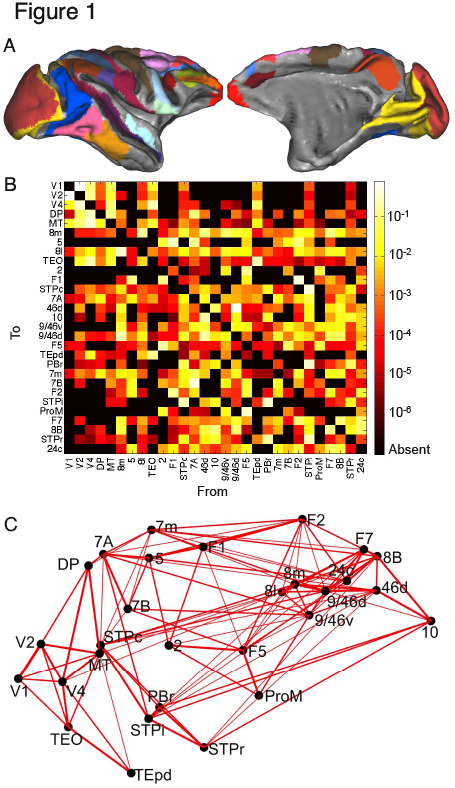
The network consists of 29 widely-distributed cortical areas. (A) Lateral (left) and medial (right) plots of the macaque cortical surface with the 29 areas marked in color. Plots generated using Caret (Van Essen et al., 2001). (B) Connection strengths between all 29 areas. The strength of the projection from area A to area B is measured by the Fraction of Labeled Neurons (FLN), defined as the number of neurons (labeled by the tracer) projecting from area A to area B divided by the total number of neurons projecting to area B from all other areas (see Experimental Procedures for more details). (C) Three dimensional positions of areas along with strongest connections between them (*FLN >* 0.005). Connection strength is indicated by line width.

Second, the dynamics of each area were described by a threshold-linear recurrent network consisting of interacting excitatory and inhibitory neural populations. The parameters of this network were calibrated by the neurophysiology of the primary visual cortex (Binzegger et al., 2009), but rescaled as described in the next paragraph. This is a highly simplified description of the overall dynamics of a cortical area and ignores most within-area variability. Nevertheless, it allows us to parsimoniously capture the essential requirements for a hierarchy of timescales. In Fig. 6 we extend our results to a model where areas show a wider variety of dynamics, and in the Discussion we suggest further extensions.

Third, we hypothesized that the local microcircuit is qualitatively canonical (Douglas and Martin, 1991), i.e. the same across areas, but that there are quantitative inter-areal differences, and that these quantitative differences play a crucial role in generating the characteristic timescales of areas. Specifically, it has been observed that the number of spines on the basal dendrites of layer 3 pyramidal neurons increases sharply from primary sensory areas to prefrontal areas (Elston, 2000; Elston et al., 2011). Taking spine count as a proxy for the number of excitatory synapses per pyramidal cell, we introduced a gradient of excitatory input strength across the cortex. We modeled this by scaling the strength of excitatory projections in an area according to the area’s position in an anatomically-derived hierarchy, which we describe below.

### Gradient of excitation strength along the cortical hierarchy

The laminar pattern of inter-areal projections can be used to place cortical areas in a hierarchy (Felleman and Van Essen, 1991; Barbas and Rempel-Clower, 1997). Such a hierarchy is defined by the observation that neurons mediating feedforward connections from one area to another tend to originate in the supragranular layers of the source area, whereas feedback projections tend to originate from infragranular layers. This observation was subsequently quantified by Barone et al. (2000), who observed that a hierarchical distance between a source area and a target area could be defined in terms of the fraction of projecting neurons located in the supragranular layers of the source area; they were then able to reproduce the hierarchy of Felleman and Van Essen (1991) using observations of connections to only two areas (V1 and V4).

Figure 2A shows the hierarchical distance measured this way for all pairs of cortical areas included in the model. We follow the approach of Markov et al. (2014b), and use these values to estimate each area’s position in an underlying hierarchy. We found that the position of an area in this anatomical hierarchy is strongly correlated with measurements of the number of spines on pyramidal neurons in that area (Elston, 2007). This allowed us to introduce a systematic gradient of excitatory connection strength per neuron along the cortical hierarchy in the model, and to explore how such cortical heterogeneity interacts with the pattern of long-range projections to produce large-scale dynamics.

**Figure 2.**
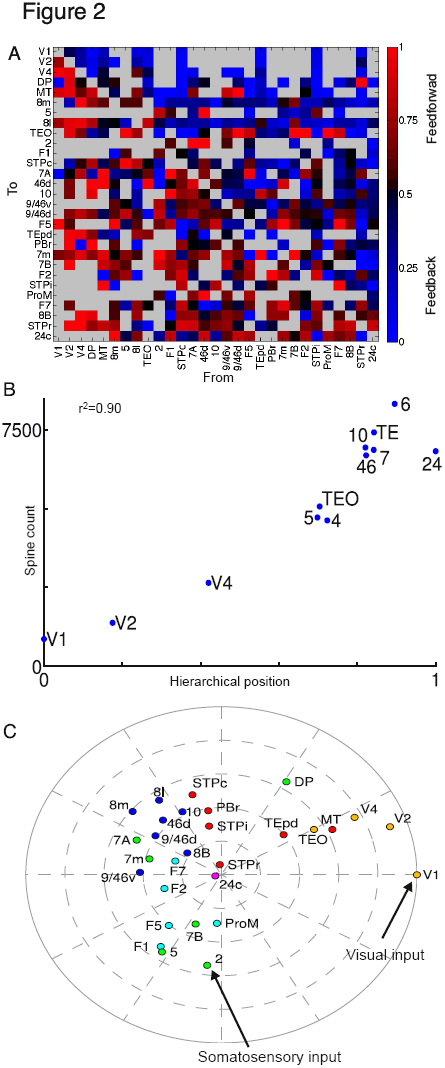
Hierarchical organization of the cortex. (A) Fraction of neurons in a projection originating from the supragranular layers of the source area (Supragranular Layer Neurons or SLN). Areas are arranged by hierarchical position. Thus most feedforward projections (SLN>0.5) lie below the main diagonal and most feedback projections (SLN <0.5) lie above the diagonal. Absent projections are shown in grey. (B) Hierarchical position of an area is well-correlated with the number of spines on pyramidal neurons in that area (Elston, 2007). For details on area labels in this panel see Supplemental Experimental Procedures. (C) Two-dimensional plot of areas determined by long-range connectivity and hierarchy. The distance of an area from the edge corresponds to its hierarchical position. The angular distance between two areas is inversely related to the strength of connection between them (i.e. closer areas are more strongly connected). Areas are colored by cortical lobe.

As a visual and conceptual aid, in Fig. 2C we use a two-dimensional circular embedding to plot hierarchy and inter-areal connectivity for the 29 areas in our model. Here the angle between two areas reflects how strongly they are interconnected (closer areas have stronger connections), and the distance of an area from the circumference reflects its position in the hierarchy (with higher areas closer to the center). The connectivity is only approximately described by this low-dimensional embedding (see Experimental Procedures), but it captures broad features of cortical organization and provides intuitive understanding of the model’s behavior. In particular, the embedding suggests two hierarchical streams of sensory input originating in area V1 (primary visual cortex) and area 2 (part of primary somatosensory cortex) respectively, and converging on a set of densely-connected association areas. We next explored the response of the network to these sensory inputs.

### Response to visual inputs

We simulated the response of the network to a pulsed input to primary visual cortex (area V1). The resulting cortical response is propagated up the visual hierarchy, progressively slowing as it proceeds (Figure 3A). Early visual areas, such as V1 and V4, exhibit fast, short-lived responses. Prefrontal areas, on the other hand, exhibit slower responses and longer integration times, with traces of the stimulus persisting several seconds after stimulation. As with the response to a pulse of input, white-noise input is integrated with a hierarchy of timescales: the activity of early sensory areas shows rapid decay of autocorrelation with time whereas cognitive areas are correlated across longer periods (Figs. 3B and C). Thus, a hierarchy of widely disparate temporal windows or timescales emerges from this anatomically-calibrated model system.

**Figure 3.**
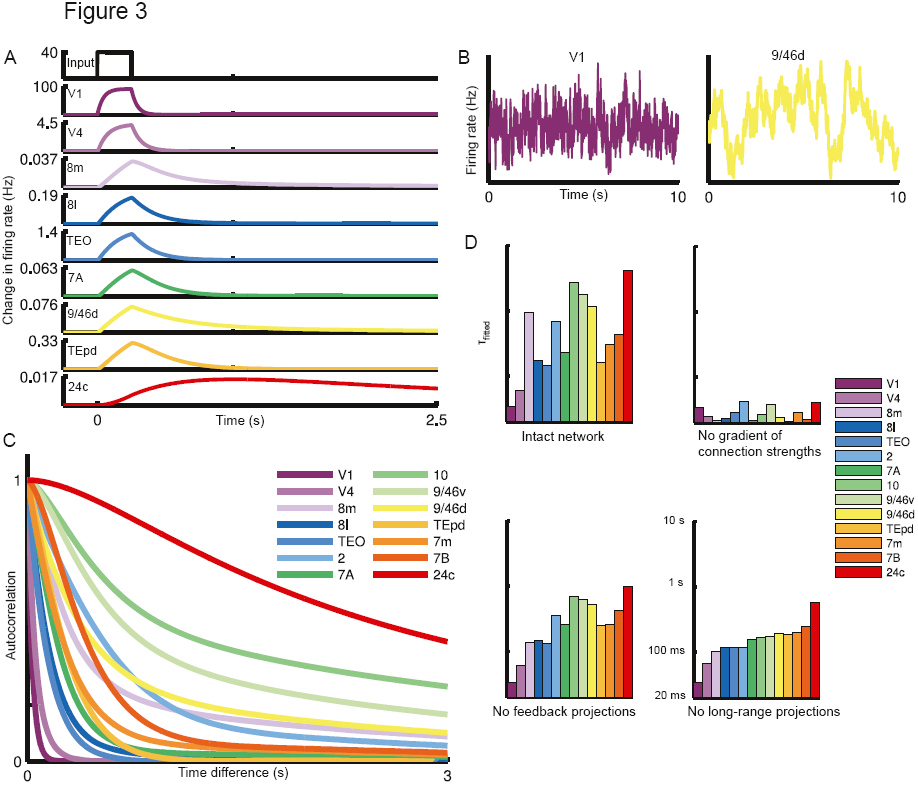
The network shows a hierarchy of timescales in response to visual input. (A) A pulse of input to area V1 is propagated along the hierarchy, displaying increasingly longer decay times as it reaches areas higher in the hierarchy. In all panels, areas are arranged by (and colored by) their position in the anatomical hierarchy. (B) Sample traces contrasting the activity of area V1 and dorsolateral prefrontal cortex in response to white-noise input to area V1. (C) Autocorrelation of area activity in response to white-noise input to V1. The autocorrelation decays with different time constants in different areas, showing a functional hierarchy ranging from area V1 at the bottom to prefrontal areas at the top. (D) The dominant time constants in various areas of the network, extracted by fitting the autocorrelation with a single or double exponential function. Top left: time constants for intact network. Time constants tend to increase along the hierarchy but depend on the influence of long-range projections. Thus, area 8m has a low hierarchical position but shows a slow timescale resulting from strong connections to prefrontal areas, while area TEpd has a high hierarchical position but is weakly connected to other slow areas and thus shows a faster timescale. Top-right: network with no gradient of excitatory synapses across areas. Bottom-left: network with feedback projections lesioned. Bottom-right: network with all long-range projections lesioned.

To quantitatively compare areas, we fit single or double exponentials to the decay of each area’s autocorrelation function (see Figure S2 for plots of the fits). These exponential fits capture a dominant characteristic timescale for each area in our model in response to visual stimulation. The time constants from the fits are plotted in the top-left panel of Figure 3D, with areas ordered by position in the anatomical hierarchy. As can be seen from the bar plot, the dominant timescale of an area tends to increase along the hierarchy (i.e. left to right), suggesting an important role for a gradient of excitation in generating the temporal hierarchy. Nevertheless, an area’s timescales are not entirely determined by its hierarchical position, and the plotted timescales do not increase monotonically with hierarchy.

Thus, the timescales of each area in our model reflect the combined effect of intrinsic hierarchical position and long-range projections. To dissect the contributions of local and long-range projections, we simulated the network after removing either inter-areal projections or differences in local microcircuitry, and fit time constants to the resulting activity. In the upper-right panel of Figure 3D, we show that the range of timescales is drastically reduced in the absence of area-specific differences in the local microcircuit. Moreover, there is no longer a relationship to an area’s position in the anatomical hierarchy. Thus, while differences in long-range inputs and outputs to each area are significant, they are not sufficient to account for disparate timescales on their own and local heterogeneity is needed.

In the lower left panel of Figure 3D, we show the effect of removing long-range feedback projections, and for the lower right panel we remove all long-range projections and simulate the activity of individual areas separately. The range of time-constants is lower, reflecting the propensity of slow areas to form long-range excitatory loops with each other. More significantly, once long-range projections are removed an area’s time-constant simply reflects its position in the hierarchy.

To gain some intuition for the role of long-range projections in the model, consider area 8m (part of the frontal eye fields), which is low in the hierarchy and would show a rapid decay of correlation in the absence of long-range projections (lower-right panel of Figure 3D). However, as can be seen from Figure 2B, it participates in a strongly-connected core of prefrontal and association areas (Ercsey-Ravasz et al., 2013; Markov et al., 2013b), and these connections allow it to show long timescales that emerge from inter-areal excitatory loops (note that these timescales are strongly attenuated in the absence of feedback projections, as in the lower-left panel of Figure 3D). The shared slower timescales are particularly characteristic of prefrontal areas in our model (see Figure S2, especially those areas that are best fit by two timescales). Conversely, while area TEpd is high in the hierarchy it does not participate in this core and is instead strongly-coupled to ventral stream visual areas. Thus, it reflects the faster timescales of visual input. In Figure S6, we extend these observations and show that scrambling the long-range projections generically reduces both the diversity and the range of temporal responses (also see Discussion).

### Multiple functional hierarchies

The response to visual input reveals an ascending hierarchy of timescales in the visual system. We next stimulated primary somatosensory cortex (area 2), which is weakly connected to the visual hierarchy and strongly connected to other somatosensory and motor areas (this organization can be seen in Figure 2C). As previously, the input propagates up a hierarchy of timescales (Figure 4A). However, the somatosensory response uncovers a different dynamical hierarchy from visual stimulation. Here, primary somatosensory cortex shows the fastest timescale, followed by primary motor cortex (area F1) and somatosensory association cortex (area 5). Parietal and premotor areas show intermediate timescales and, as in response to visual stimulation, prefrontal areas show long timescales (Figure 4A).

**Figure 4.**
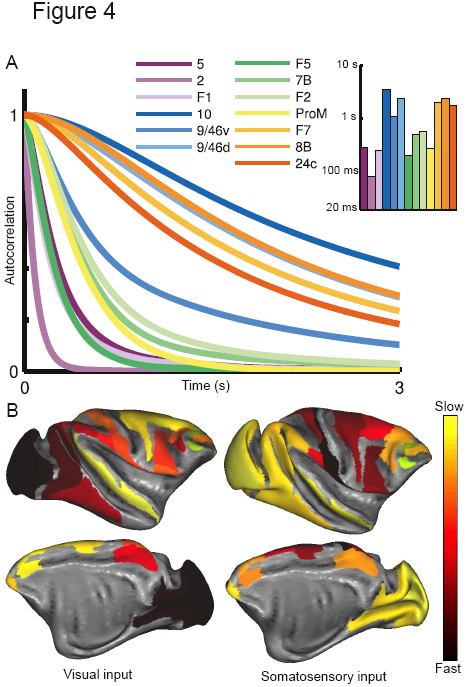
The response to somatosensory input reveals a different functional hierarchy subserved by the same anatomical network. (A) Autocorrelation of activity for areas that show strong responses to white-noise input to area 2 (part of primary somatosensory cortex). Area labels are arranged according to position in the underlying anatomical hierarchy. Inset: time constants fitted to the autocorrelation function for each area. (B) Timescales in response to visual (left) and somatosensory input (right) shown with lateral (top) and medial (bottom) views of the cortex.

In Figure 4B, we contrast time-constants in response to visual and somatosensory stimulation by plotting them on a map of the cortical surface. Visual areas demonstrate much weaker responses than before, and are mostly driven by top-down projections from association areas. Thus, in the absence of direct input they reflect the slower timescales of a distributed network state.

An area’s timescales emerge from a combination of local circuit properties, the specificity of long-range projections, and the particular input to the network. Our model allows us to examine the contribution of each. These can be intuitively summarized by noting that each area in Figure 2C shows timescales approximately determined by its distance from the periphery (hierarchical position and amount of recurrent excitation), its proximity to the central clusters (long-range connectivity) and its distance from the source of input.

### Localized eigenvectors and separated timescales

The model we used for a single area is threshold-linear, meaning that we ignore nonlinearities besides the constraint that firing rates remain positive. This allowed us to explore the genesis of separated timescales using the tools of linear system analysis. The activity of a linear network is the weighted sum of a set of characteristic activity patterns, called eigenvectors (Rugh, 1995). Each eigenvector evolves on a different timescale, given by the corresponding eigenvalue. The input determines the degree to which an eigenvector is driven. Thus the eigenvectors set the space of possible responses of the system, and the inputs determine which response is actually observed.

We found that the eigenvectors of the linearized network are localized, meaning that each eigenvector is concentrated around a small region of the network. As can be seen in Figure 5, the eigenvectors corresponding to short timescales are broadly concentrated around sensory areas and the eigenvectors corresponding to long timescales are concentrated at frontal areas. In general, if an eigenvector is small at a particular node then its amplitude at that node in response to input will also be small, and the corresponding timescale will be only weakly expressed. Here, the localization of eigenvectors means that, for most inputs, the network dynamics will be dominated by rapid timescales at sensory areas and slower timescales at cognitive areas. In previous theoretical work, we showed how such localized eigenvectors could arise in networks with gradients of local properties, and investigated their relationship to a diversity of timescales (Chaudhuri et al., 2014).

**Figure 5.**
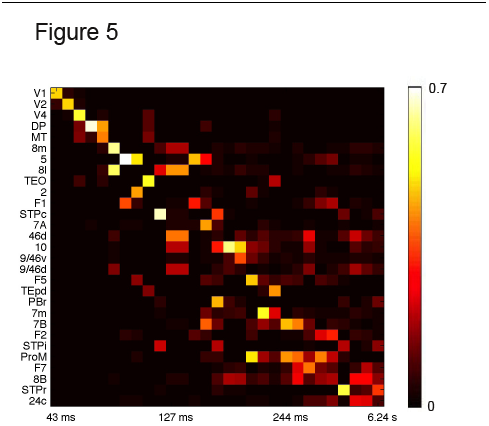
Eigenvectors of the network coupling matrix are weakly localized, corresponding to segregated temporal modes. Each column shows the amplitude of an eigenvector at the 29 areas, with corresponding timescale labeled below. The 29 slowest eigenvectors of the system are shown.

### Extension to a system with nonlinear dynamics and multistability

In our network we have used a threshold-linear model for local areas. The simplicity of this local circuit model allowed us to highlight the essential requirements for a hierarchy of timescales and to provide intuition for the model behavior from linear systems theory. Moreover, many systems can be linearly approximated around a state of interest, and neural responses are often near-linear over a wide range of inputs (Wang, 1998; Chance et al., 2002). This makes linear and threshold-linear dynamical equations useful models for neural circuits (Dayan and Abbott, 2001).

Nevertheless, linear models show a limited range of dynamical behaviors and, in particular, are unable to capture features such as persistent activity or multistability, which are thought to be important for cognitive capabilities in higher areas (Wang, 2013). To demonstrate how our model might be extended, we replaced our local circuit model with a firing rate (“mean-field”) version of a spiking network with more realistic synaptic dynamics (Wang, 2002; Wong and Wang, 2006). As shown in Figure 6A, when isolated a local area in this network can display qualitatively different dynamical regimes. When recurrent connections are relatively weak, an area shows a single stable state. As the strength of recurrent excitation is increased, the area shows a transition to a regime with two stable states, distinguished by low and high firing rates and corresponding to a resting state and a self-sustained persistent activity state. In this regime, an area can integrate inputs over time and can show persistent activity in the absence of a stimulus. Such dynamical regimes have been proposed as a general model for “cognitive-type” computations such as working memory and decision-making (Wang, 2002; 2013).

**Figure 6.**
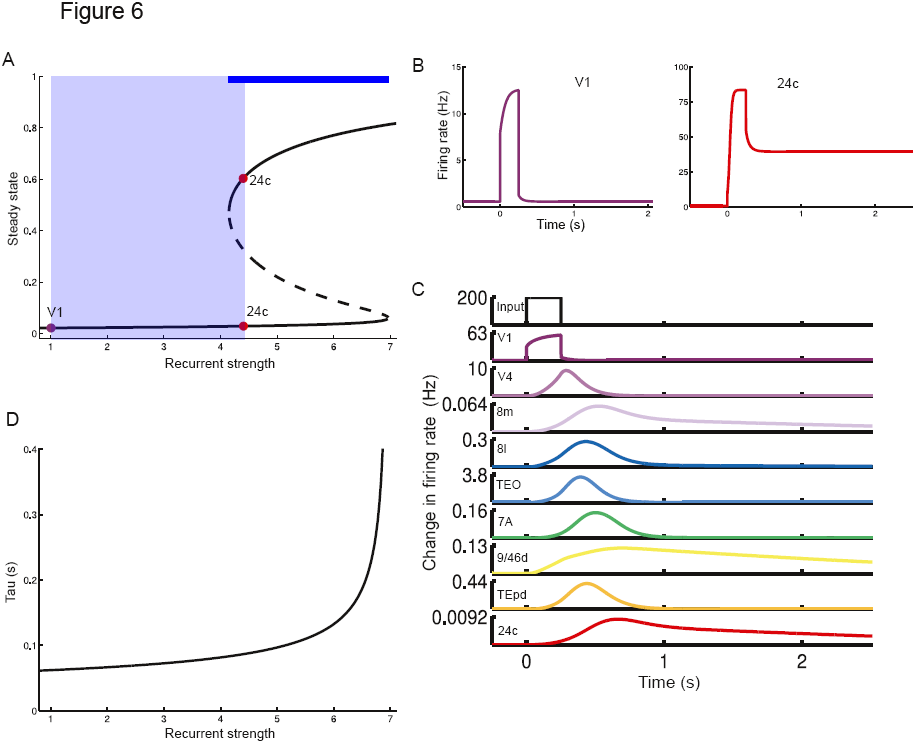
Hierarchy of timescales in a nonlinear model. (A) Possible steady-states (bifurcation diagram) for an area in the network as a function of recurrent strength (normalized by the value at V1). Stable steady-states are shown with solid lines. Areas with comparatively low recurrent strength (such as V1) display only a single steady-state. Increasing the recurrent connection strengths leads to a regime with a high-activity steady-state (such as seen in area 24c). The dashed line indicates an unstable steady-state lying intermediate to the stable states. The thick blue line above the plot indicates the parameter regime within which bistability exists, while the light blue shaded region indicates the range of recurrent strengths used for areas in the model. Steady-states are shown as fractional activation of NMDA conductance (see Experimental Procedures for more details). (B) Response of disconnected areas to a strong pulse of input, shown for two areas at either end of the hierarchy. As in (A), V1 is only able to show a single stable state, while area 24c shows sustained delay activity. (C) The timescales of responses to a small perturbation serve as a probe of the recurrent strength of a local area. These timescales are much smaller than those shown in response to a larger input, but they emerge from the same underlying gradient in recurrent strengths. (**D**) Response of connected network to a brief pulse of input to area V1. As in Figure 3, the input is propagated up the network, slowing down as it proceeds along the hierarchy. Note that the input is not strong enough to switch any area into the high-activity stable state.

Using this model for each local area in the large-scale network, we introduced the same gradient as before for the excitation strength in an area as a function of its position in the hierarchy (Figure 2). Consequently, early sensory areas show single stable states while areas at the top of the hierarchy can also show persistent activity when driven by strong inputs, which shift the steady-state of the network (Figure 6B). Smaller perturbations are insufficient to shift the state of a node; nevertheless such perturbations take longer to decay away in areas further up the hierarchy (Figure 6C).

In response to small inputs the network shows similar behavior to our threshold-linear model: the response to a brief input to area V1 is propagated up the cortical hierarchy, with rapid decays in early sensory areas and slow decays in association areas (Figure 6D). Thus, the results obtained previously extend to a nonlinear attractor model with a larger dynamical repertoire. While exploring the more complex dynamical behaviors that this network can show is beyond the scope of the current paper, one interesting consequence of the extended model is that the timescales of small fluctuations around baseline predict the ability of an area to show much longer timescales in response to larger inputs (Figure 6C, and see Discussion), as observed in Honey et al. (2012) and Murray et al. (2014).

### Functional connectivity

Thus far we have focused on the response properties of individual areas. We now investigate the implications of local circuit heterogeneity and a hierarchy of timescales for the organization of the network as measured by correlations in resting-state activity (resting-state functional connectivity). In our model, frontal and association areas reflect a slowly varying state of the network, and we hypothesized that this network state should strongly shape functional connectivity.

In Figure 7A we computed functional connectivity in our threshold-linear model with heterogeneity in local area properties, or without it (as typically assumed in previous models relating functional to anatomical connectivity). The inclusion of a gradient of local excitation reduced the correlation (*r*^2^) between functional and anatomical connectivity from 0.83 to 0.53 (in Figure S7 we show similar results after convolution with a BOLD kernel (Boynton et al., 1996)).

**Figure 7.**
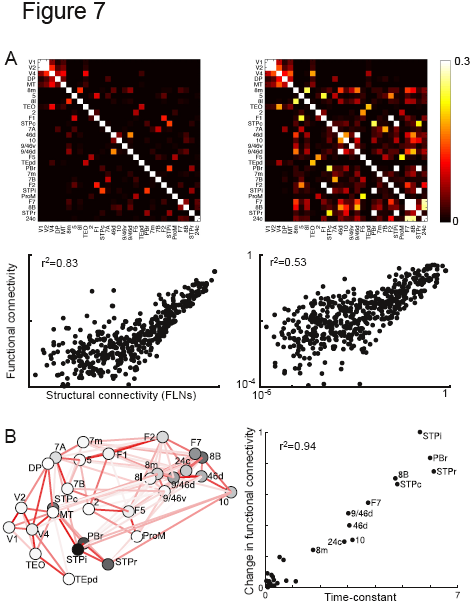
Functional connectivity depends on local microcircuitry. (**A**) Functional connectivity for two networks with identical long-range connectivity. The network on the left has the same local properties at each area, while the network on the right has a gradient of local recurrent strengths. Top panel: correlations between area activity in response to background white noise input to each area. Bottom panel: functional connectivity (correlations) vs. structural connectivity (FLNs) for non-zero projections. The network with a gradient of local recurrent strengths shows enhanced functional connectivity for slow areas, and a smaller overall correlation between functional and anatomical connectivity (showing that long-range connections alone cannot predict the global brain activity patterns). (**B**) Effect of lesioning areas, one at a time, on functional connectivity. Left panel: Darker areas are those with a greater influence on resting-state functional connectivity, as measured by the normalized changes in functional connectivity after lesioning. Right panel: The effect of lesioning an area on functional connectivity is well-correlated with the time constant of spontaneous fluctuations in that area. See Figure S5 for the time-constants of resting-state activity for each area and Figure S7 for activity after convolution with a BOLD kernel.

A number of studies have tried to connect long-range anatomical connectivity to correlations in resting-state activity and have found that the presence and strength of an anatomical connection between areas partially predicts correlations in neurophysiological signals from those areas, but there are significant differences (Hagmann et al., 2008; Honey et al., 2009; Damoiseaux and Greicius, 2009; Honey et al., 2010; Deco and Corbetta, 2011; Deco et al., 2014). Our results also suggest that inter-areal connections alone are insufficient to predict the system’s activity pattern. Importantly, we find that heterogeneity in local anatomical connectivity could help account for the previously unexplained variance.

In our model we find that slower frontal and temporal areas in particular show enhanced functional connectivity. Consequently, areas exhibiting slower timescales play a predominant role in shaping the network, as explicitly shown by the effects of lesioning individual areas (Figure 7B, left panel). In the simple case of identical input to each area, the effect of lesioning an area is well predicted by the time constant of intrinsic fluctuations (Figure 7B, right panel). Note that the areas most important for functional connectivity are not simply those at the highest positions in the hierarchy, and we find that hierarchy alone poorly predicts impact on functional connectivity. For instance, the caudal superior temporal polysensory region (STPc) and the rostral parabelt (PBr) are at intermediate hierarchical positions, but they have strong connections to other parts of the superior temporal polysensory region (strong connections shown by the darker lines in Figure 7B) forming a cluster that shapes functional connectivity. In general, cortical areas endowed with a combination of an intermediate to high hierarchical position and strong connections to other slow areas exhibit the strongest influence on the global activity patterns.

## Discussion

The main findings of the present work are threefold. First and foremost, it establishes a circuit mechanism for the emergence of a hierarchy of temporal receptive windows, which has received empirical support in recent human experiments (Hasson et al., 2008; Lerner et al., 2011; Honey et al., 2012; Gauthier et al., 2012) and single-unit monkey experiments (Murray et al., 2014). This model extends working memory from local circuits (Wang, 2001) to a large-scale brain system across multiple timescales. Second, it demonstrates that conventional graph-theoretical and network analyses of functional connectivity must be revised: inter-areal heterogeneity implies that areas cannot be treated as identical nodes of a network and slow dynamics in the prefrontal and other association areas can play a disproportionate role in determining the statistical pattern of functional connectivity. And third, this is the first large-scale dynamical model of the macaque monkey cortex based on weighted and directed connectivity and that takes into account heterogeneity across cortical areas.

The ability to integrate and internally hold information across time is critical for higher functions such as decision-making. Accumulation of information in a decision process involves neural integrators with a long time constant (Smith and Ratcliff, 2004; Gold and Shadlen, 2007; Wang, 2008; Brunton et al., 2013). On the other hand, the brain must be able to rapidly and transiently respond to constantly changing external stimuli with a short time constant. Complex behavior thus requires the coexistence of a multitude of timescales, ranging from fast timescales of sensory responses to slow timescales of integrative cognitive processes. We demonstrate how such a hierarchy of timescales (or temporal receptive windows) naturally emerges in a large-scale model of primate cortex, calibrated by quantitative anatomical data.

While the observed temporal hierarchy depends on areas having different recurrent connection strengths, inter-areal connections play a significant role and we show that the range of time constants and their distribution are dramatically reduced in the absence of long-distance connections (Figure 3D, lower panels compared with control in upper left panel). Moreover, the specificity of long-range connections has an important role to play. As demonstrated in Figure S6A, scrambling the long-range connections while retaining differences in local properties makes areas show similar time-constants to each other; this effect is especially prominent for fast visual areas. Preserving the topology of the network (i.e., which area is connected to which) but scrambling the weights has a similar, though smaller effect (Figure S6B). Thus, both the pattern of connections and their weights help separate time constants.

Our work revealed multiple functional hierarchies converging on a network of frontal and other association areas, as seen anatomically in Figure 2C and functionally in the timescales of Figures 3 and 4 and the correlation matrices of Figure 7. In our data the prefrontal and association areas preferentially participate in densely-connected subnetworks, forming a “core” (Markov et al., 2013b; Ercsey-Ravasz et al., 2013; van den Heuvel et al., 2012), and sensory inputs from multiple streams enter this slow distributed network to be integrated and potentially fed back to early sensory areas. This large-scale organization is reminiscent of global workspace cognitive architectures (Dehaene et al., 1998).

In order to examine timescales in the clearest way possible, the present work used a threshold-linear rate model, which summarizes the effective dynamics of local areas. In this model, time constants are well defined (mathematically as the inverse of the real parts of eigenvalues of a linear system). However, we also explicitly showed that the main results hold when the local circuit model is highly nonlinear, endowed with multiple attractor states (multi-stability). It is also noteworthy that this work did not focus on the latency of neural response (Schmolesky et al., 1998; Bullier, 2001), for which a spiking neural model is needed. Comparing the model with neurophysiological data, one may wonder if neurons in the monkey cortex actually display slow responses during stimulus presentation as shown in the model. In fact, in decision tasks neurons of the prefrontal and parietal cortices do show quasi-linear ramping activity over a long period of time (Smith and Ratcliff, 2004; Gold and Shadlen, 2007; Wang, 2008), with a time constant that in some cases may appear to be effectively infinite (Brunton et al., 2013). Therefore, our model is the simplest that is adequately designed to reveal a hierarchy of timescales in the primate cortex.

To systematically introduce heterogeneity across all the cortical areas in our model, we assigned each area to a position according to its hierarchical rank, which was determined by its pattern of feedforward and feedback projections (Markov et al., 2014b). A priori, there is no reason why excitatory input would vary systematically along the anatomically defined hierarchy. However, as shown in Figure 2B, we find the hierarchical position of a cortical area correlates very strongly with the number of spines per neuron in that area. This suggests an underlying organizational principle in the cortex, which could be explored in future analyses (see also Scholtens et al. (2014) for a similar observation and Barbas and Rempel-Clower (1997) and Hilgetag et al. (2002) for evidence that hierarchical position correlates with the degree of lamination and relative density of a cortical area).

While there are no systematic measurements of the characteristic timescales of multiple cortical areas in response to different stimuli, a number of recent studies have compared temporal responses and integration timescales across cortical areas and report a hierarchical organization (Hasson et al., 2008; Ogawa and Komatsu, 2010; Lerner et al., 2011; Bernacchia et al., 2011; Honey et al., 2012; Gauthier et al., 2012; Murray et al., 2014). In particular, the study of Honey et al. (2012) connected a functional hierarchy in the timescales of preferred stimuli to a dynamical hierarchy evidenced by the timescales of correlation in network activity, and found autocorrelation timescales broadly similar to those we model (in particular, see Figure 6 of Honey et al. (2012)). Similarly, Murray et al. (2014) found that autocorrelation traces were well-described by exponentials, the hierarchical ordering of areas they observe agrees with our model, and the timescales of small fluctuations in their study are close to the intrinsic time-constants of areas in the model (i.e., in the absence of long-range projections such as in Fig. 3D, lower right panel).

Our model has several testable predictions. Though there are multiple combinations of local time constants and network connection strengths that could give rise to a particular set of observed network time constants, the model suggests that timescales of small fluctuations should reflect the intrinsic time constants of areas (lower right panel of Figure 3D), while those of larger, task-dependent responses should reflect time constants that emerge from the entire system (upper left panel of Figure 3D). In the model, the slow network timescales are driven by strongly-connected frontal and temporal areas (Markov et al., 2013a;b), corresponding to a slowly varying state of the network. Inactivating these areas should decrease slow dynamics observed in connected areas lower in the hierarchy. The differential responses to visual and somatosensory input suggest that when a particular input is not involved in a task, the corresponding sensory areas better reflect slow changes in global cortical state. This may explain decreases in low-frequency power (i.e. slow modes) seen in ECoG recordings when a subject engages in a task (He et al., 2010). Finally, we predict that areas with longer timescales, such as prefrontal and superior temporal areas, can shape functional connectivity to a relatively greater degree. This highlights the importance of incorporating heterogeneous local dynamics of cortical areas in studying the determinants of functional connectivity and, intriguingly, suggests that functional connectivity might be used to probe local properties. While there is some evidence that frontal and association areas show generally enhanced functional connectivity (Sepulcre et al., 2010) as well as of a correlation between enhanced functional connectivity and slow timescales (Baria et al., 2013), it would be interesting to use functional imaging to better understand the link between functional connectivity and timescales of response (for example, as determined using the approaches of Hasson et al. (2008); Lerner et al. (2011); Honey et al. (2012); Gauthier et al. (2012)).

Although most of the results were obtained with a threshold-linear model for local areas, the same hierarchy of timescales holds when local areas were modeled by a highly nonlinear attractor network (Figure 6). The microcircuit we use here is a simplified variant of one previously proposed as a model for general “cognitive-type” computations, like working memory and decision-making (Wang, 2002; 2013). While we do not explore the broader range of dynamical behaviors in this network, note that in this extended model the timescales of small fluctuations around baseline predict the ability of an area to show much longer timescales in response to larger inputs. This can be seen by comparing the timescales of Figure 6C with the steady-states of Figure 6A, and by contrasting responses to large and small perturbations in Figures 6B and D (also note that the timescales in response to large perturbations tend to be slower than those in response to small perturbations even if the area does not show bistability). This suggests a possible explanation for the observations of Honey et al. (2012), who find that the timescales of spontaneous fluctuations in an area (with time constants on the order of hundreds of milliseconds) correlate with its sensitivity to temporal structure in stimuli lasting multiple seconds. Similar observations were also made in Murray et al. (2014), who found that the timescales of spontaneous fluctuations were correlated with slower baseline activity and with the timescales of neurons encoding reward across trials.

Another interesting feature of this model architecture is that areas can show persistent activity emerging either locally or from inter-areal loops; these may have different functional roles to play and should be explored in future work.

Our model is parsimonious with several simplifications, designed to capture a basic mechanism for the generation of a hierarchy of timescales. The model can be improved and extended in several ways in future work. First, in our model activity propagates along the cortical hierarchy with significant attenuation, which might be solved by implementing more sophisticated feedback projections onto inhibitory as well as excitatory neurons (Moldakarimov et al., 2015). Second, it will be interesting to incorporate a laminar structure in the model. Third, we only consider cortico-cortical connections. While these connections form the major input to a cortical area (Markov et al., 2011), subcortical projections may play an important role in setting cortical state and gating information flow. Fourth, we used two global parameters to scale long-range connection strengths. While this seems a reasonable first step, it is only an approximation (Shepherd et al., 2005) and as data relating anatomy and physiology continues to emerge (Mao et al., 2011) it should be incorporated into a future iteration of the model. Fifth and finally, extensions of our model should include other heterogeneities observed between areas, such as in interneuron subtypes and their densities (Medalla and Barbas, 2009) and in neuromodulatory signaling (Hawrylycz et al., 2012).

In conclusion, we report a novel, quantitatively-calibrated, dynamical model of the macaque cortex with directed and weighted connectivity. The identification of a specific circuit mechanism for a hierarchy of timescales (temporal receptive windows) represents a key advance towards understanding specialized processes and functions of different (from early sensory to cognitive-type) cortical areas. Our findings demonstrate the importance of heterogeneity in local areal properties, as well as the specific profile of long-range connectivity, in sculpting the large-scale dynamical organization of the brain.

## Experimental Procedures

### Anatomical data

The connectivity data are from an on-going project to quantitatively measure all connections between cortical areas in the macaque brain, with areas defined according to a 91 area parcellation scheme (Markov et al., 2014a). Briefly, connection strengths between areas are measured by counting the number of neurons labeled by retrograde tracer injections. The number of neurons labeled in a projection ranges from a few neurons to on the order of 100,000 neurons. In order to control for injection size, these counts are then normalized by the total number of neurons labeled in the injection, yielding a fractional weight or FLN (Fraction of Labeled Neurons) for each pathway, defined as

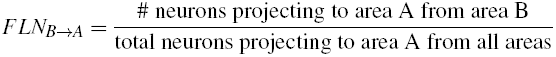

The corresponding weights span 5 orders of magnitude. So far, 29 areas have been injected and we use the subnetwork consisting of these areas. In this network the presence or absence of all connections is known bidirectionally, and 66% of possible connections exist in the network, though with widely varying strengths.

We also use data on the fraction of neurons in each projection that originate in the upper layers of the source area (called the SLN, for Supragranular Layer Neurons (Markov et al., 2014b)), defined as

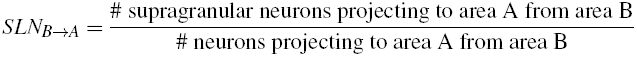

### Hierarchy and the low-dimensional connectivity embedding

To extract the hierarchy, we follow observations from the visual system that the fraction of projections originating in the supragranular layers of the source area (the SLN) serves as a measure of hierarchical distance between the source and target areas (Felleman and Van Essen, 1991; Barone et al., 2000; Markov et al., 2014b). We then use a generalized linear model to assign hierarchical values to areas such that the differences in hierarchical values predict the SLNs (similar to the method used in Markov et al. (2014b)).

For the two-dimensional circular embedding of Figure 2C, we use − log(FLN(*A*_*i*_, *A*_*j*_)) as a measure of dissimilarity between areas *A*_*i*_ and *A _j_* and compute angles *θ*_*i*_ such that the angular distances between areas correspond to the dissimilarity. We fix area V1 to have *θ* = 0 (choosing another area to fix rotates the plot). Finally, we plot the areas on a 2-dimensional polar plot with *θ* (*A*_*i*_) = *θ*_*i*_ and 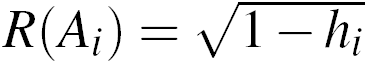.

See the Supplemental Experimental Procedures and Figure S1 for an expanded discussion of the hierarchy and the circular embedding.

### Model architecture

Each area consists of an excitatory and an inhibitory population described by

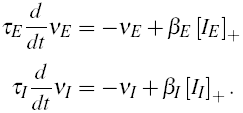

*ν*_*E*_ is the firing rate of the excitatory population, with intrinsic time constant *τ*_*E*_ and input current *I*_*E*_, and for which the f-I curve has slope *β*_*E*_. [*I*_*E*_ ]+ = max(*I*_*E*_, 0). The inhibitory population has corresponding parameters *τ*_*I*_, *I*_*I*_ and *β*_*I*_. Values for *τ*_*E*_, *τ*_*I*_, *β*_*E*_ and *β*_*I*_ are taken from Binzegger et al. (2009).

At each area, the input currents have a component from within the area (i.e. local input) and another that comes from other areas:

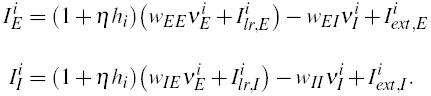

*w*_*EE*_ and *w*_*EI*_ are couplings to the excitatory population from the excitatory and inhibitory population respectively, 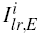 is the long-range input to the excitatory population, and 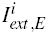 is external input (both stimulus input and any noise we add to the system). *w*_*IE*_, *w*_*II*_, 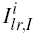 and 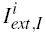 are corresponding parameters for the inhibitory population.

We scale the excitatory inputs to an area, both local and long-range, by its position in the hierarchy, *h*_*i*_. Note that *h*_*i*_ is normalized between 0 and 1, and *η* is a scaling parameter that controls the effect of hierarchy. By setting *η* = 0 we remove intrinsic differences between areas.

The values of the local parameters are: *τ*_*E*_ =20 ms, *τ*_*I*_=10 ms, *β*_*E*_ =0.066 Hz/pA, *β*_*I*_=0.351 Hz/pA, *w*_*EE*_ = 24.3 pA/Hz, *w*_*EI*_ = 19.7 pA/Hz, *w*_*IE*_ = 12.2 pA/Hz and *w*_*II*_ = 12.5 pA/Hz. For more details on how we choose these parameters see Supplemental Experimental Procedures.

Long-range input is modeled as excitatory current to both excitatory and inhibitory cells:

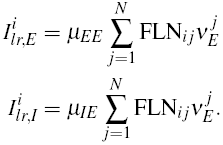

Here *j* ranges over all areas. 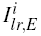 and 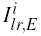 are the inputs to the excitatory and inhibitory populations, 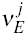 is the firing rate of the excitatory population in area *j* and FLN_*ij*_ is the FLN from area *j* to area *i*. *μ*_*EE*_ and *μ*_*IE*_ are scaling parameters that control the strengths of long-range input to the excitatory and inhibitory populations, respectively, and do not vary between connections; all the specificity comes from the FLNs. The network thus has three parameters: *μ*_*EE*_ and *μ*_*IE*_ control the connection strengths of long-range projections, and *η* maps the hierarchy into excitatory connection strengths. We set *μ*_*EE*_ = 33.7 pA/Hz, *μ*_*IE*_ = 25.3 pA/Hz, and *η* = 0.68.

Note that we ignore inter-areal conduction delays in most of our simulations, but see Figure S3 for a network with conduction delays added in.

### Pulse input, autocorrelation and fitted time constants

For Figures 3, 4 and 7, we choose the background input for each area so that the excitatory population has a firing rate of 10 Hz and the inhibitory population has a rate of 35 Hz.

In Figure 3A, V1 receives a 250 ms pulse of input that drives its firing rate to 100 Hz. For the remaining panels, the stimulus to V1 is white-noise with a mean of 2 Hz and a standard deviation of 0.5 Hz. The other areas receive a small amount of background input (standard deviation on the order of 10−^5^), but are primarily driven by long-range input propagating out from area V1. For Figure 4, the currents are the same except that area 2 receives the stimulus rather than V1.

For each area we extract time constants by fitting both one and two exponentials to the part of the autocorrelation function that decays from 1 to 0.05. If the sum of squared errors of the single exponential fit is less than 8 times that of the double exponential, then we report the time-constant of the single exponential. Otherwise, we use the sum of the time-constants from the double exponential fit, with each weighted by its corresponding amplitude. The fits and time constants in response to V1 and area 2 input and for resting state activity are shown in Figures S2, S4 and S5.

For Figure 4B, we map the time-constants logarithmically to a heatmap and plot them using Caret (Van Essen et al., 2001).

### Functional connectivity

The simulations shown in Figure 7A highlight the effect of intrinsic hierarchy on functional connectivity. Here, we contrast a network without hierarchy with a network that has a gradient of local excitatory connections but, unlike the network shown in the remaining figures, no gradient in the long-range projection strengths (thus the two networks have exactly the same long-range connection strengths and any differences emerge purely from local properties). The input currents become

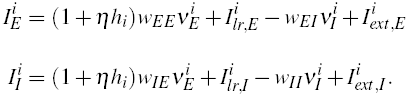

For Figure 7B we use the same network as in Figures 3 and 4, so that all incoming excitatory projections to an area are scaled by the area’s position in the hierarchy.

We calculate functional connectivity as the correlation matrix of area activity in response to equal white-noise input to all areas. For Figure 7B we use the analytical solution for the correlation matrix (see Supplemental Experimental Procedures) to speed up calculations. The change in functional connectivity upon lesioning a given area, A, is measured as ‖*C*_*l,A*_ − *C*_*rs,A*_‖/ ‖*C*_*rs,A*_‖, where *C*_*l,A*_ is the correlation matrix of the network after lesioning area A, *C*_*rs,A*_ is the correlation matrix of the intact network with the row and column corresponding to A removed, and the double lines indicate the norm of the matrix. These values are then scaled to lie between 0 and 1. The timescales are fit to the activity of the intact network, using the same single and double exponential fits described previously.

### Nonlinear network

The single area model for the nonlinear network is a variant of a model proposed in Wong and Wang (2006) as an mean-field approximation to a spiking neural network with AMPA, GABA and NMDA synapses (Wang, 2002). Each area is described by the equations:

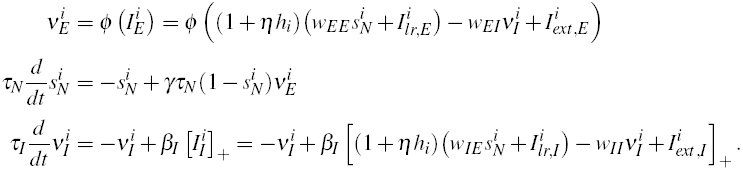

*ν*_*E*_ and *ν*_*I*_ are excitatory and inhibitory firing rates, *s*_*N*_ is a gating variable corresponding to the activity of NMDA synapses (with decay timescale *τ*_*N*_) and *ϕ* is a simplified f-I curve from Abbott and Chance (2005). We elaborate on these equations in the Supplemental Experimental Procedures. Note that *s*_*N*_ is bounded between 0 and 1: in the differential equation for *s*_*N*_ the input is multiplied by 1 − *s*_*N*_, which becomes small as *s*_*N*_ approaches 1. This leads to a response that saturates for high currents and allows the system to have two stable states in certain regimes. Parameter values are: *τ*_*N*_ = 60 ms, *τ*_*I*_ = 10 ms, *γ* = 0.641, *w*_*EE*_ = 250.2 pA, *w*_*EI*_ = 8.110 pA/Hz, *w*_*IE*_ = 303.9 pA and *w*_*II*_ = 12.5 pA/Hz.

For Figure 6A-C we set the long-range connections to 0 and characterize an isolated area. The bifurcation diagram of Figure 6A is generated by plotting the steady-states of the network as we vary the hierarchy parameter (written as 1 + *ηh_i_* in the equation above). For each parameter value, we also calculate the Jacobian matrix about the low firing rate steady-state (i.e. we linearize about that steady-state) and in Figure 6C we plot the timescale of the slowest eigenmode of this linearized system.

For Figure 6B we set *η* = 3.4 and give a 100 Hz pulse of input for 250 ms to the two disconnected areas at opposite ends of the hierarchy (V1 and 24c) and plot the response as each area decays to steady-state. For Figure 6D we consider a connected network, with long-range projections only targeting excitatory subpopulations, for simplicity. Thus 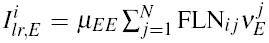, and we set *μ*_*EE*_ = 0.5. We give a 200 Hz pulse of input to area V1 for 250 ms and plot the response for selected areas.

## Supplemental Figure Legends

**Figure S1.**
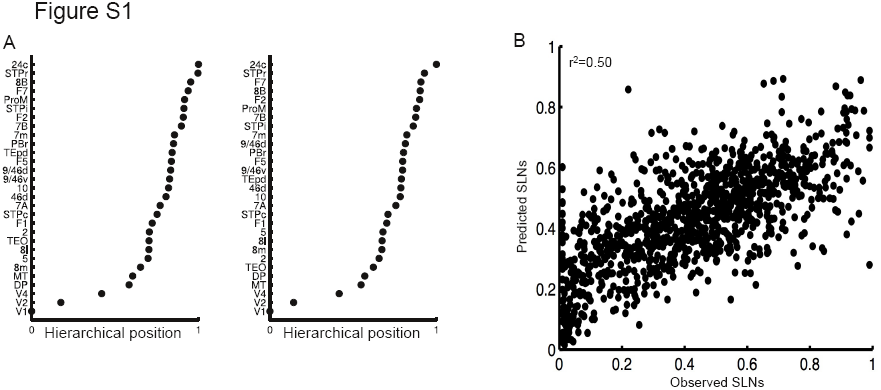
Hierarchy fitted from pairwise SLN relationships. (A) Left panel: Hierarchy fitted from logistic regression (and used in main text). The hierarchical position of an area is normalized to lie between 0 and 1. Right panel: Hierarchy fitted from beta regression (Cribari-Neto and Zeileis, 2010). (B) SLN values predicted from logistic regression compared to observed SLNs.

**Figure S2.**
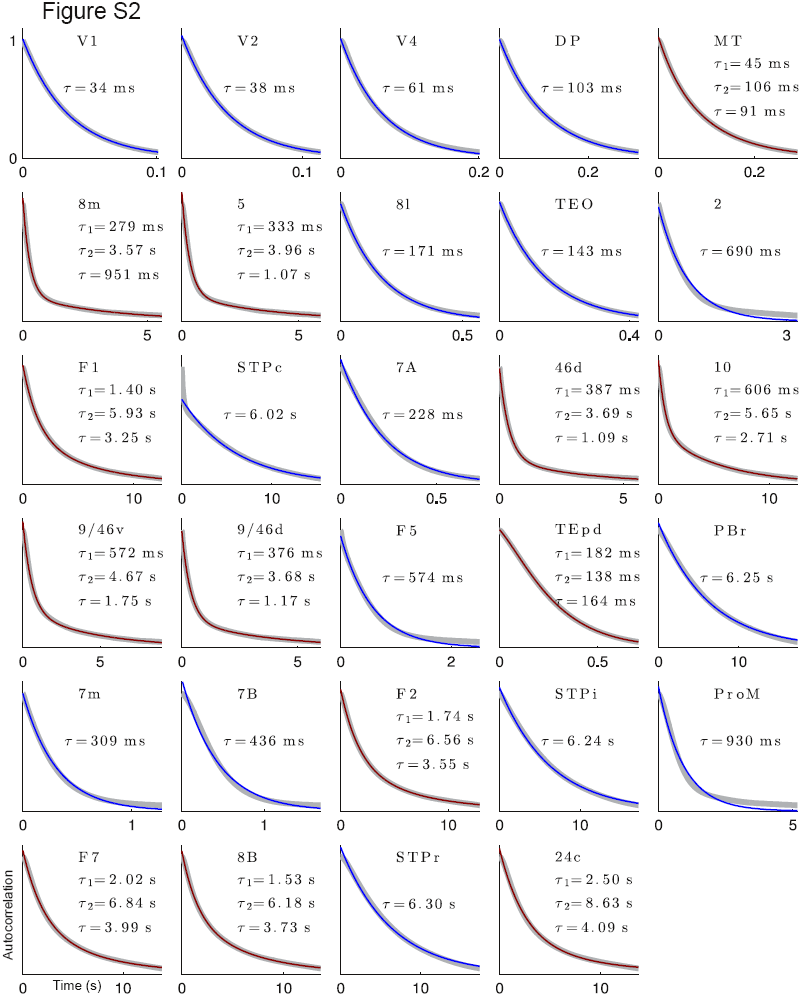
Timescales in response to white-noise input to V1. Data shown in grey, single exponential fits in blue and double exponential fits in dark red. For double exponential fits, *τ*_1_ and *τ*_2_ are the time-constants of individual exponentials, and *τ* is a weighted average of *τ*_1_ and *τ*_2_, with weights given by the amplitudes of the exponentials.

**Figure S3.**
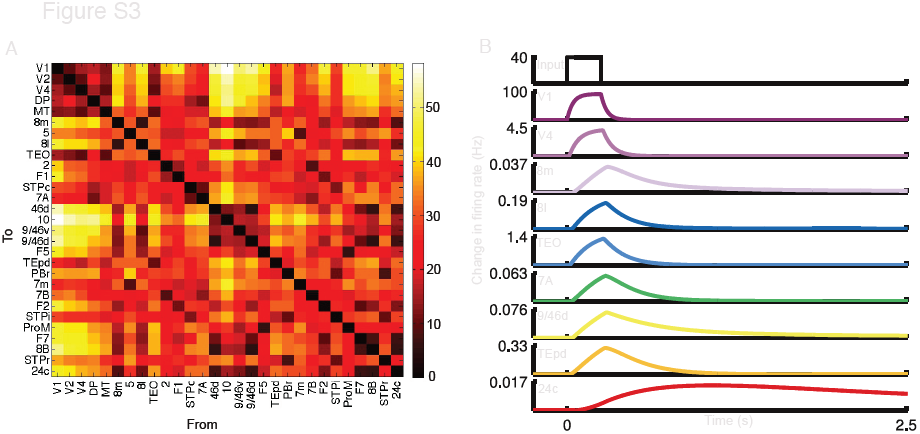
Response of a network with inter-areal conduction delays. (A) Distances (in mm) between the nodes of the network (Ercsey-Ravasz et al., 2013). (B) Response of the network to a pulse of input to area V1. Conduction delays between nodes are imposed using the distances in panel A and a conduction velocity of 1.5 m/s (Deco et al., 2009).

**Figure S4.**
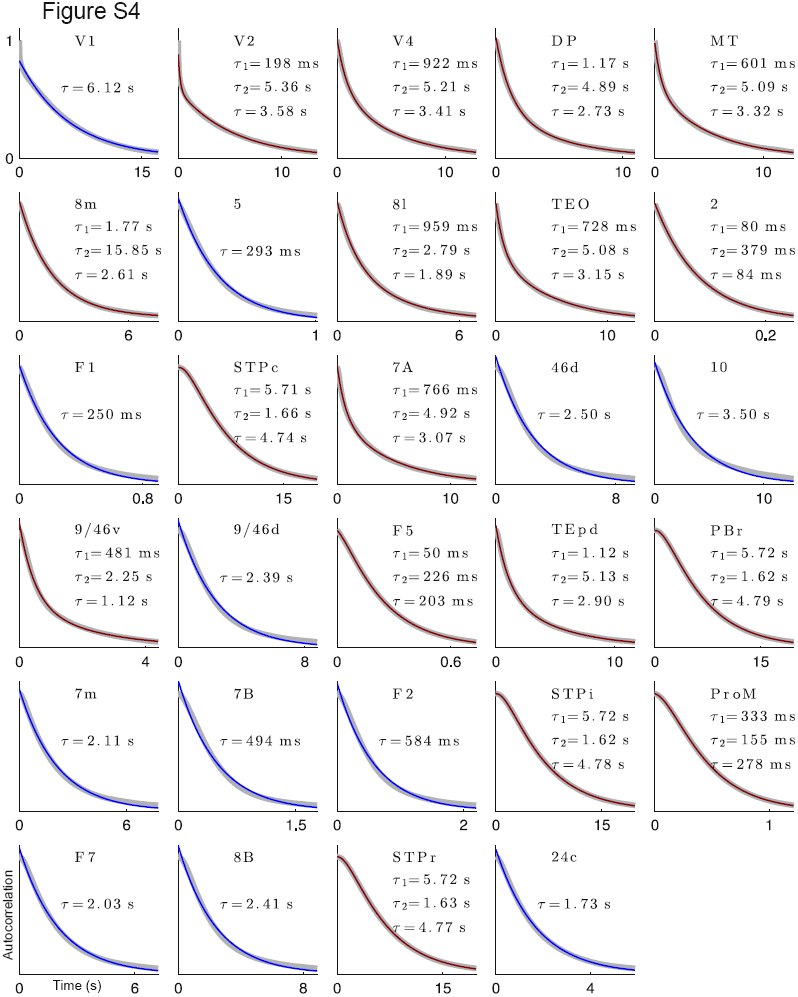
Timescales from exponential fits of activity in response to white-noise input to Area 2. Colors as in Figure S2.

**Figure S5.**
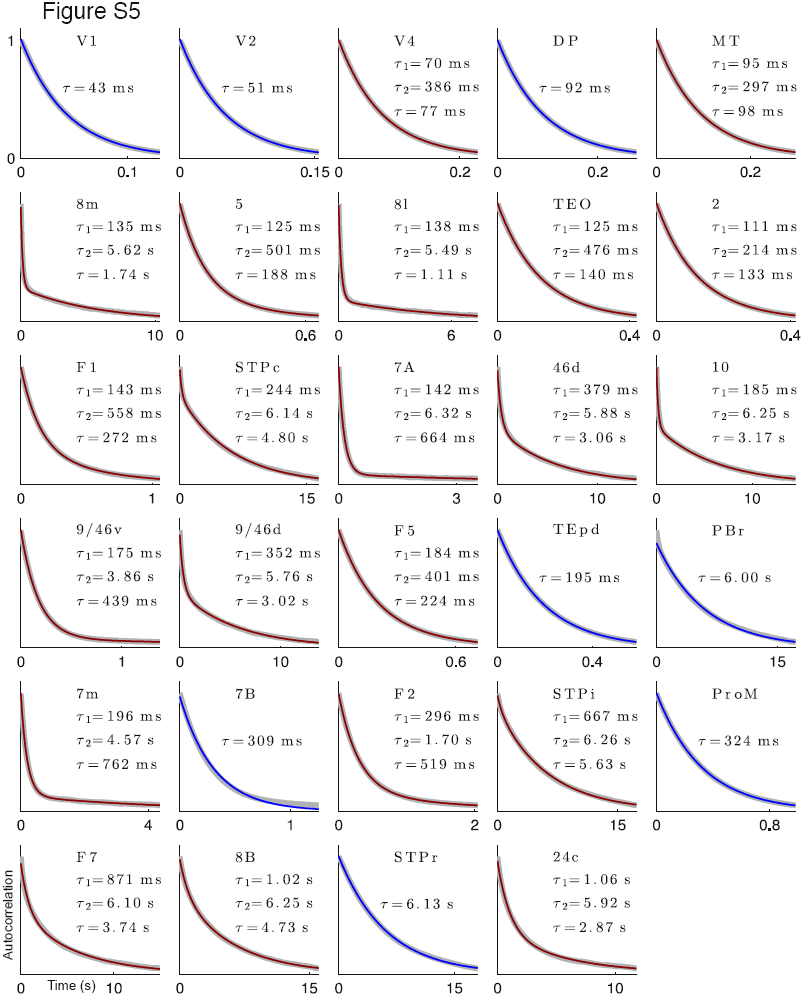
Timescales from exponential fits of resting-state activity (i.e., equal white-noise input to all areas). Colors as in Figure S2.

**Figure S6.**
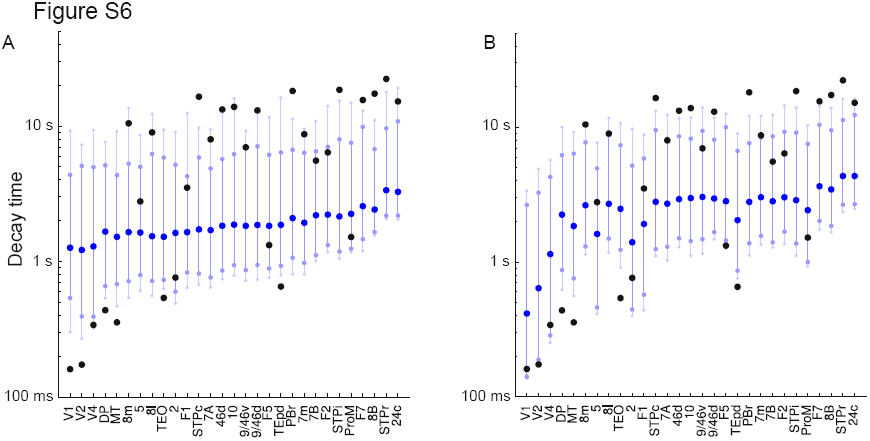
Effect of scrambling long-range connectivity on resting-state network dynamics, measured by the time taken for an area’s activity to return to within 5% of baseline after a 250 ms pulse of input. (A) Distribution of timescales when all connection strengths are randomly permuted. Dark blue circles indicate median value, lighter blue circles mark the 10th and 90th percentiles, and the very light blue circles mark 5th and 95th percentiles. Values for intact network are shown in black, for comparison. Areas in scrambled networks are much more similar to each other (compare black to blue), and fast visual areas show the greatest disruption. (B) Distributions when only non-zero connection strengths are permuted, thus preserving the connectivity pattern but not strengths. As before, median shown in dark blue, 10% − 90% range in lighter blue, and 5% − 95% in very light blue, along with intact times in black.

**Figure S7.**
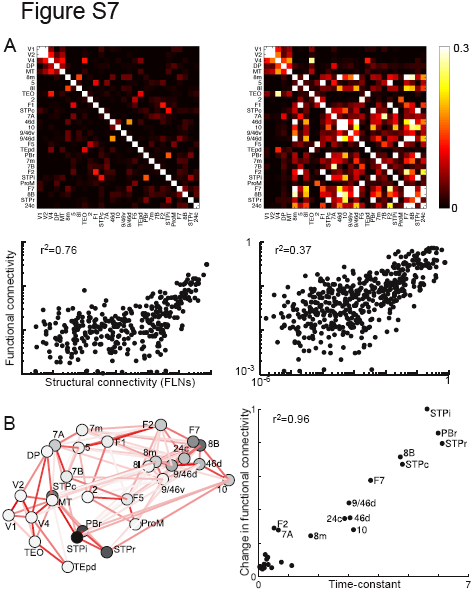
Functional connectivity of simulated BOLD signal. (A) As in Figure 7A, the network on the left has the same local properties at each node, while the network on the right has a gradient of local recurrent strengths. Firing rate is convolved with a gamma function to generate a simulated BOLD signal (Boynton et al., 1996). Top panel: functional connectivity in response to background white noise input to each node. Bottom panel: functional connectivity (correlations in BOLD) vs. structural connectivity (FLN) for non-zero projections. (B) Effect of lesioning areas on functional connectivity measured via simulated BOLD signal. Plots are as in Figure 7B.

## Supplemental Experimental Procedures

Several of these sections are expanded versions of the corresponding sections in Experimental Procedures in the main text. To make these descriptions self-contained, the relevant portions from the main text are repeated here.

### Hierarchy and low-dimensional connectivity embedding

In the visual system, projections directed from early visual areas to higher-order areas (i.e. increasing size of receptive field, position-invariance, and so on) tend to originate in the supragranular layers of the cortex and terminate in layer 4 (Felleman and Van Essen, 1991; Barone et al., 2000). Conversely, projections from higher-order areas to early visual areas originate in the infragranular layers and terminate outside of layer 4. This observation was systematized by Felleman and Van Essen (1991), who used these anatomical constraints to place cortical areas in a hierarchical ordering.

Felleman and Van Essen used a discrete classification of projections: in their framework projections are either feedforward, feedback or lateral depending on where the majority of projections originate and terminate. However, such binary relations are typically insufficient to specify a unique hierarchy (Hilgetag et al., 1996). Subsequently, it was observed that rather than classifying a projection as feedforward, feedback or lateral, the fraction of neurons in a projection originating in the supragranular layers (the SLN) could be used as a continuous measure of hierarchical displacement: the difference of the SLN from 50% is positive for feedforward projections and negative for feedback projections, and its magnitude gets larger as a projection moves further away from lateral (Barone et al., 2000). For example, a projection with an SLN of 90% would be very strongly feedforward, while a projection with an SLN of 65% would be only moderately feedforward. Using these values, the Felleman and van Essen hierarchy could be reproduced using observations of connections to only two areas (V1 and V4) (Barone et al., 2000).

To construct the hierarchy we follow a similar framework to Markov et al. (2014b) and use a generalized linear model. We assign hierarchical values to each area such that the difference in values predicts the SLN of a projection. Specifically, we assign a value *H*_*i*_ to each area *A*_*i*_ such that

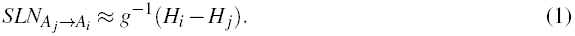

We choose *g*−1 to be a logistic function (logistic regression), which is standard for probabilities and fractional values, but we note that other functions yield similar values (Figure S1A). We have one such constraint for each projection (536 in total), and we find the set of hierarchical values that best fit these constraints. In the fit we weight the contribution of each projection by the log of its FLN to preferentially match stronger and less noisy projections. The resulting best fit hierarchy is shown in the left panel of Figure S1A. We then normalize by the maximum hierarchical value yielding *h*_*i*_ = *H*_*i*_/*H*_*max*_.

We extract the spine counts in Figure 2B from Elston (2007) and plot the areas in common with our data set. The parcellation in that paper is coarser than the parcellation we use, so we report the results in terms of that parcellation. For area 7 we average together the hierarchical positions of 7A, 7B and 7m; for 6 we average F2, F5 and F7; and for 46 we average together 46d, 9/46d and 9/46v.

For the two-dimensional circular embedding of Figure 2C, we convert the FLN to a measure of dissimilarity according to

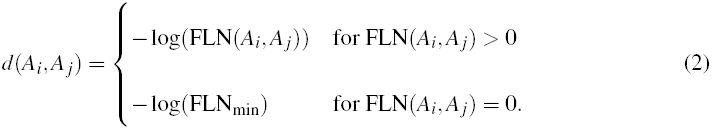

Here, *A*_*i*_ is the *i*th area, and FLN_min_ is some value less than the smallest FLN in the network. We use FLN_min_ = 10^−7^ but the results are robust to the precise choice of this value. We then assign angles *θ*_*i*_ to each area such that *d*(*A*_*i*_, *A*_*j*_) ≈ *R* min(*|θ_i_ − θ _j_|,* 2*π − |θ_i_ − θ _j_*|), where *R* is a single free parameter. We fix area V1 to have *θ* = 0, but choosing any other area to fix would simply rotate the plot. Finally, we plot the areas on a 2-dimensional polar plot with *θ*(*A*_*i*_) = *θ*_*i*_ and 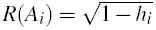

### Model architecture

Each of the 29 nodes consists of an excitatory and an inhibitory population, which summarize the effective dynamics of the area. Populations are described by

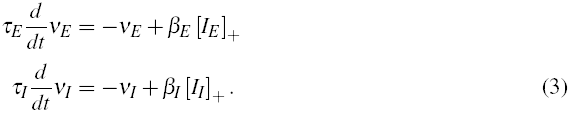

*ν*_*E*_ is the firing rate of the excitatory population, with intrinsic time constant *τ*_*E*_ and input current *I*_*E*_, and for which the f-I curve has slope *β*_*E*_. [*I*_*E*_ ]+ = max(*I*_*E*_, 0). The inhibitory population has corresponding parameters *τ*_*I*_, *I*_*I*_ and *β*_*I*_. Values for *τ*_*E*_, *τ*_*I*_, *β*_*E*_ and *β*_*I*_ are given below and are taken from Binzegger et al. (2009).

At each node, the input currents have a component from within the area (i.e. local input) and another that comes from other areas:

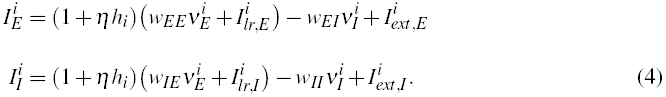

*w*_*EE*_ and *w*_*EI*_ are couplings to the excitatory population from the excitatory and inhibitory population respectively,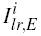 is the long-range input to the excitatory population, and 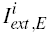 is external input (both stimulus input and any noise we add to the system). *w*_*IE*_, *w*_*II*_, 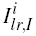 and 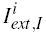 are corresponding parameters for the inhibitory population.

Following Binzegger et al. (2009), we write *w*_*ij*_ = *α*_*i*_*S*_*ij*_, where *i* and *j* can be E or I. *α*_*E*_ (*α*_*I*_) measures charge introduced per excitatory (inhibitory) spike times transmitter release probability; both are slightly modified from Binzegger et al. (2009). *S*_*ij*_ is the number of synapses from cells of type *j* to cells of type *i*, taken from the counts for layer 2/3 cells in Binzegger et al. (2004). Inhibitory values are weighted averages of basket, double bouquet and chandelier cells, with weights chosen according to their projections to the excitatory population.

We scale the excitatory inputs to an area, both local and long-range, by its position in the hierarchy, *h*_*i*_. *h*_*i*_ is normalized between 0 and 1, and *η* is a scaling parameter that controls the effect of hierarchy. By setting *η* = 0 we remove intrinsic differences between areas. Note that we scale both local and long-range projections with hierarchy, rather than just local projections, in accordance with the observations of Markov et al. (2011), who find that the proportion of local to long-range connections is approximately conserved across areas.

The values of the local parameters are: *τ*_*E*_ =20 ms, *τ*_*I*_=10 ms, *β*_*E*_ =0.066 Hz/pA, *β*_*I*_=0.351 Hz/pA, *α*_*E*_ = 0.007 pC, *α*_*I*_ = 0.025 pC, *w*_*EE*_ = 24.3 pA/Hz, *w*_*EI*_ = 19.7 pA/Hz, *w*_*IE*_ = 12.2 pA/Hz and *w*_*II*_ = 12.5 pA/Hz.

Long-range input is modeled as excitatory current to both excitatory and inhibitory cells:

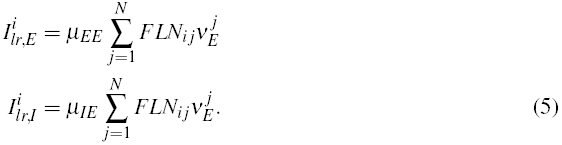

Here *j* ranges over all areas. 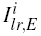 and 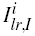 are the inputs to the excitatory and inhibitory populations, 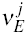 is the firing rate of the excitatory population in area *j* and *FLN*_*ij*_ is the FLN from area *j* to area *i*. *μ*_*EE*_ and *μ*_*IE*_ are scaling parameters that control the strengths of long-range input to the excitatory and inhibitory populations, respectively, and do not vary between connections; all the specificity comes from the FLNs. The network thus has three parameters: *μ*_*EE*_ and *μ*_*IE*_ control the connection strengths of long-range projections, and *η* maps the hierarchy into excitatory connection strengths.

We can choose the excitatory to inhibitory ratio of an input current, *γ* = *I*_*inp,E*_ /*I*_*inp,I*_, such that the steady-state firing rate of the excitatory population does not change when the current is present. Given input of *I*_*inp,E*_ to the excitatory population, an input of *γI_inp,E_* to the inhibitory population increases the inhibitory firing rate sufficiently to cancel out the additional input to the excitatory population. We call such inputs balanced. We choose *μ*_*EE*_ and *μ*_*IE*_ with a ratio slightly above this value so that projections are weakly excitatory.

For our simulations, we use *μ*_*EE*_ = 33.7 pA/Hz, *μ*_*IE*_ = 25.3 pA/Hz, and *η* = 0.68.

### Network with conduction delays

In our simulations we ignore conduction delays between areas. While these will be important for oscillations, synchronization and other fine temporal structure, the timescales we consider are typically slow enough that conduction delays should not play an important role.

In Figure S3 we demonstrate that our results hold in a network with realistic conduction delays. We use distances from the same data set as the connectivity strengths (Ercsey-Ravasz et al., 2013) and, to ensure a fair comparison, assume a relatively low conduction velocity of 1.5 m/s (Deco et al., 2009). As shown in Figure S3B, the response of this network to a pulse of input to area V1 is almost identical to that of a network without conduction delays.

### Scrambled connectivity

For the simulations shown in Figure S6A, we scramble the connectivity matrix by permuting all entries of the matrix randomly. For Figure S6B, we preserve the absent entries and permute the non-zero entries. Note that the connectivity data show specificity both in terms of which projections exist and in their strengths, and both the probability of a connection and its strength decay exponentially with distance between areas (Markov et al., 2011; 2013; 2014a; Ercsey-Ravasz et al., 2013). In particular, nearby areas tend to be strongly connected and to have similar timescales (see Fig. 2C); thus scrambling projections should reduce the separation of timescales.

We examine the response of these scrambled networks to a pulse of input to all areas, similar to the “resting-state” condition. In the intact network, areas are dominated by a few timescales and are well fit by one or two summed exponentials. However, a number of the scrambled networks show responses that consist of many mixed timescales and are not well described by two exponentials. Thus we use a non-parametric measure of timescale: we compute the time taken after pulse offset for the area’s activity to decay to within 5% of its value at baseline. Scrambling the connection strengths makes about 20% of networks unstable, meaning that responses to input grow instead of decaying, and we exclude these networks. We then compute the median and the 5th, 10th, 90th and 95th percentile of the decay time distribution for each area, and contrast it with values for the intact network.

### Functional connectivity for a linear network

If a linear network is driven by white noise input then, away from the threshold, it evolves according to the equation

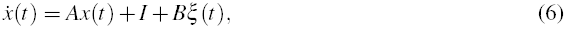

where *I* is the mean of the noise, *B* is its covariance matrix and *A* is the coupling matrix, which includes any intrinsic leak of activity.

In the steady-state the covariance, *C*, of this matrix is the solution to the equation (Gardiner, 1985)

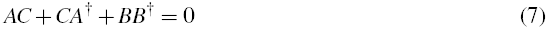

This equation can be solved given the eigenvector basis (Deco et al., 2013). In the eigenvector decomposition, *A* = *V* Λ *V* ^−1^, where Λ is the diagonal matrix of eigenvalues and the columns of *V* are the right eigenvectors of A. Define

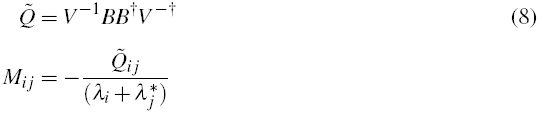

Then *C* = *V MV*^†^.

As an aid to intuition, assume that *A* is a normal matrix so that *V* ^−1^ = *V* ^†^. Then 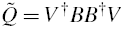, and the covariance matrix of the network is a rescaled version of the covariance structure of the input noise.

If, as in the simulations of Figure 6, the input noise is independent and identical at each node, then the covariance matrix of the noise is diagonal with constant entries (and all correlations come from the structure of the network). If this has the value *σ*^2^ at each node then, for a normal matrix, 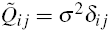, and *M* is diagonal with *i*th entry *τ*_*i*_*σ*^2^/2, where *τ*_*i*_ = −*1*/*λ*_*i*_. Hence the covariance of the *i*th eigenmode is proportional to its corresponding timescale.

Now *C* = *V MV*^†^, meaning that the matrix is rotated out of the eigenvector basis giving a non-diagonal matrix. Thus eigenvectors that are more broadly shared contribute more to the functional connectivity.

### Functional connectivity with hemodynamic response function

For Figure S6, we convolve the firing rates of the excitatory population at each node with a hemodynamic response function of the form

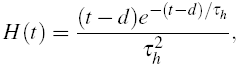

with timescale *τ*_*h*_ = 1.25 s and delay *d* = 2.25 s (Boynton et al., 1996). This yields a simulated BOLD signal, and we calculate the functional connectivity as the correlation matrix of this activity.

### Nonlinear network

The single area model is a variant of the model developed in Wong and Wang (2006) as a simplified mean-field version of the spiking network of Wang (2002). There the dynamics were assumed to be dominated by the slow time-constant of NMDA synapses, and the activity of the inhibitory population was incorporated into the effective connection strengths between the excitatory populations. As in that study, we assume that the dynamics of the excitatory population are modeled by a dimensionless gating variable, *s*_*N*_, reflecting the fractional activation of the NMDA conductance, with timescale set by the slow NMDA time-constant. However, we also consider an inhibitory population, modeled with a threshold-linear differential equation (as in the previous sections).

The equation for the excitatory population is

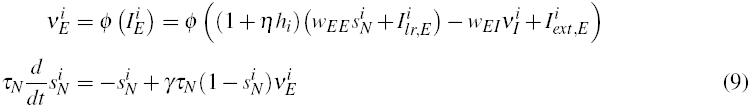

Here *ν*_*E*_ is the excitatory firing rate and *s*_*N*_ is the NMDA gating variable, which is bounded between 0 and 1. *ϕ* models the firing rate-current dependence of a leaky integrate-and-fire neuron (Abbott and Chance, 2005) and is defined as

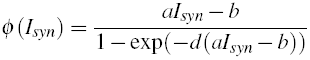

with *a* = 0.27 Hz/pA, *b* = 108 Hz and *d* = 0.154 s.

The inhibitory population is described with a threshold-linear equation as before.

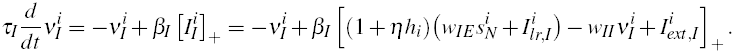

Parameter values are: *τ*_*N*_ = 60 ms, *τ*_*I*_ = 10 ms, *γ* = 0.641, *w*_*EE*_ = 250.2 pA, *w*_*EI*_ = 8.110 pA/Hz,

*w*_*IE*_ = 106.7 pA and *w*_*II*_ = 4.402 pA/Hz.

## References

Abbott L.F., and Chance F.S. (2005). Drivers and modulators from push-pull and balanced synaptic input. Prog. Brain Res. 149, 147&155.

Amit D.J., Fusi S., and Yakovlev V. (1997). Paradigmatic working memory (attractor) cell in IT cortex. Neural Comput. 9, 1071&1092.

Barbas H., and Rempel-Clower N. (1997). Cortical structure predicts the pattern of corticocortical connections. Cereb. Cortex 7, 635&646.

Baria A.T., Mansour A., Huang L., Baliki M.N., Cecchi G.A., Mesulam M.M., and Apkarian A.V. (2013). Linking human brain local activity fluctuations to structural and functional network architectures. Neuroimage 73, 144&155.

Barone P., Batardiere A., Knoblauch K., and Kennedy H. (2000). Laminar distribution of neurons in extrastriate areas projecting to visual areas V1 and V4 correlates with the hierarchical rank and indicates the operation of a distance rule. J. Neurosci. 20, 3263&3281.

Bernacchia A., Seo H., Lee D., and Wang X.J. (2011). A reservoir of time constants for memory traces in cortical neurons. Nat. Neurosci. 14, 366&372.

Binzegger T., Douglas R.J., and Martin K.A. (2009). Topology and dynamics of the canonical circuit of cat V1. Neural Netw 22, 1071&1078.

Boynton G.M., Engel S.A., Glover G.H., and Heeger D.J. (1996). Linear systems analysis of functional magnetic resonance imaging in human V1. J. Neurosci. 16, 4207&4221.

Brunton B.W., Botvinick M.M., and Brody C.D. (2013). Rats and humans can optimally accumulate evidence for decision-making. Science 340, 95&98.

Bullier J. (2001). Integrated model of visual processing. Brain Res. Rev. 36, 96&107.

Bullmore E., and Sporns O. (2009). Complex brain networks: graph theoretical analysis of structural and functional systems. Nat. Rev. Neurosci. 10, 186&198.

Chance F.S., Abbott L.F., and Reyes A.D. (2002). Gain modulation from background synaptic input. Neuron 35, 773&782.

Chaudhuri R., Bernacchia A., and Wang X.J. (2014). A diversity of localized timescales in network activity. Elife 3, e01239.

Curtis C.E., and Lee D. (2010). Beyond working memory: the role of persistent activity in decision making. Trends Cogn. Sci. (Regul. Ed.) 14, 216&222.

Damoiseaux J.S., and Greicius M.D. (2009). Greater than the sum of its parts: a review of studies combining structural connectivity and resting-state functional connectivity. Brain Struct Funct 213, 525&533.

Dayan P., and Abbott L.F. (2001). Theoretical Neuroscience (The MIT Press).

Deco G., and Corbetta M. (2011). The dynamical balance of the brain at rest. Neuroscientist 17, 107&123.

Deco G., Ponce-Alvarez A., Hagmann P., Romani G.L., Mantini D., and Corbetta M. (2014). How local excitation-inhibition ratio impacts the whole brain dynamics. J. Neurosci. 34, 7886&7898.

Dehaene S., Kerszberg M., and Changeux J.P. (1998). A neuronal model of a global workspace in effortful cognitive tasks. Proc. Natl. Acad. Sci. U.S.A. 95, 14529&14534.

Douglas R.J., and Martin K.A. (1991). A functional microcircuit for cat visual cortex. J. Physiol. (Lond.) 440, 735&769.

Elston G.N. (2000). Pyramidal cells of the frontal lobe: all the more spinous to think with. J. Neurosci. 20, RC95.

Elston G.N. (2007). Specialization of the neocortical pyramidal cell during primate evolution. In Evolution of Nervous Systems: A Comprehensive Reference, J.H. Kass, and T.M. Preuss, eds. (New York: Elsevier), vol. 4, pp. 191&242.

Elston G.N., Benavides-Piccione R., Elston A., Manger P.R., and Defelipe J. (2011). Pyramidal cells in prefrontal cortex of primates: marked differences in neuronal structure among species. Front Neuroanat 5, 2.

Ercsey-Ravasz M., Markov N.T., Lamy C., Van Essen D.C., Knoblauch K., Toroczkai Z., and Kennedy H. (2013). A predictive network model of cerebral cortical connectivity based on a distance rule. Neuron 80, 184&197.

Felleman D.J., and Van Essen D.C. (1991). Distributed hierarchical processing in the primate cerebral cortex. Cereb. Cortex 1, 1&47.

Gauthier B., Eger E., Hesselmann G., Giraud A.L., and Kleinschmidt A. (2012). Temporal Tuning Properties along the Human Ventral Visual Stream. J. Neurosci. 32, 14433&14441.

Ghosh A., Rho Y., McIntosh A.R., Kotter R., and Jirsa V.K. (2008). Noise during rest enables the exploration of the brain’s dynamic repertoire. PLoS Comput. Biol. 4, e1000196.

Gold J.I., and Shadlen M.N. (2007). The neural basis of decision making. Annu. Rev. Neurosci. 30, 535&574.

Hagmann P., Cammoun L., Gigandet X., Meuli R., Honey C.J., Wedeen V.J., and Sporns O. (2008). Mapping the structural core of human cerebral cortex. PLoS Biol. 6, e159.

Hasson U., Yang E., Vallines I., Heeger D.J., and Rubin N. (2008). A hierarchy of temporal receptive windows in human cortex. J. Neurosci. 28, 2539&2550.

Hawrylycz M.J., Lein E.S., Guillozet-Bongaarts A.L., Shen E.H., Ng L., Miller J.A., van de Lagemaat L.N., Smith K.A., Ebbert A., Riley Z.L., et al. (2012). An anatomically comprehensive atlas of the adult human brain transcriptome. Nature 489, 391&399.

He B.J., Zempel J.M., Snyder A.Z., and Raichle M.E. (2010). The temporal structures and functional significance of scale-free brain activity. Neuron 66, 353&369.

Hilgetag C.C., Dombrowski S.M., and Barbas H. (2002). Classes and gradients of prefrontal cortical organization in the primate. Neurocomputing 44, 823&829.

Histed M.H., Pasupathy A., and Miller E.K. (2009). Learning substrates in the primate prefrontal cortex and striatum: sustained activity related to successful actions. Neuron 63, 244&253.

Honey C.J., Kotter R., Breakspear M., and Sporns O. (2007). Network structure of cerebral cortex shapes functional connectivity on multiple time scales. Proc. Natl. Acad. Sci. U.S.A. 104, 10240&10245.

Honey C.J., Sporns O., Cammoun L., Gigandet X., Thiran J.P., Meuli R., and Hagmann P. (2009). Predicting human resting-state functional connectivity from structural connectivity. Proc. Natl. Acad. Sci. U.S.A. 106, 2035&2040.

Honey C.J., Thesen T., Donner T.H., Silbert L.J., Carlson C.E., Devinsky O., Doyle W.K., Rubin N., Heeger D.J., and Hasson U. (2012). Slow cortical dynamics and the accumulation of information over long timescales. Neuron 76, 423&434.

Honey C.J., Thivierge J.P., and Sporns O. (2010). Can structure predict function in the human brain? Neuroimage 52, 766&776.

Hubel D.H. (1988). Eye, Brain, and Vision: Scientific American Library Series (Scientific American Press, New York).

Hubel D.H., and Wiesel T.N. (1962). Receptive fields, binocular interaction and functional architecture in the cat’s visual cortex. J. Physiol. (Lond.) 160, 106&154.

Kennedy H., Knoblauch K., and Toroczkai Z. (2013). Why data coherence and quality is critical for understanding interareal cortical networks. Neuroimage 80, 37&45.

Kiebel S.J., Daunizeau J., and Friston K.J. (2008). A hierarchy of time-scales and the brain. PLoS Comput. Biol. 4, e1000209.

Kobatake E., and Tanaka K. (1994). Neuronal selectivities to complex object features in the ventral visual pathway of the macaque cerebral cortex. J. Neurophysiol. 71, 856&867.

Lerner Y., Honey C.J., Silbert L.J., and Hasson U. (2011). Topographic mapping of a hierarchy of temporal receptive windows using a narrated story. J. Neurosci. 31, 2906&2915.

Mao T., Kusefoglu D., Hooks B.M., Huber D., Petreanu L., and Svoboda K. (2011). Long-range neuronal circuits underlying the interaction between sensory and motor cortex. Neuron 72, 111&123.

Markov N.T., Ercsey-Ravasz M., Lamy C., Ribeiro Gomes A.R., Magrou L., Misery P., Giroud P., Barone P., Dehay C., Toroczkai Z., Knoblauch K., Van Essen D.C., and Kennedy H. (2013a). The role of long-range connections on the specificity of the macaque interareal cortical network. Proc. Natl. Acad. Sci. U.S.A. 110, 5187&5192.

Markov N.T., Ercsey-Ravasz M., Van Essen D.C., Knoblauch K., Toroczkai Z., and Kennedy H. (2013b). Cortical high-density counterstream architectures. Science 342, 1238406.

Markov N.T., Ercsey-Ravasz M.M., Ribeiro Gomes A.R., Lamy C., Magrou L., Vezoli J., Misery P., Falchier A., Quilodran R., Gariel M.A., et al (2014a). A weighted and directed interareal connectivity matrix for macaque cerebral cortex. Cereb. Cortex 24, 17&36.

Markov N.T., Misery P., Falchier A., Lamy C., Vezoli J., Quilodran R., Gariel M.A., Giroud P., Ercsey-Ravasz M., Pilaz L.J., et al. (2011). Weight consistency specifies regularities of macaque cortical networks. Cereb. Cortex 21, 1254–1272.

Markov N.T., Vezoli J., Chameux P., Falchier A., Quilodran R., Huis-soud C., Lamy C., Misery P., Giroud P., Ullman S., et al (2014b). The anatomy of hierarchy: feedforward and feedback pathways in macaque visual cortex. J. Comp. Neurol. 522, 225&259.

Medalla M., and Barbas H. (2009). Synapses with inhibitory neurons differentiate anterior cingulate from dorsolateral prefrontal pathways associated with cognitive control. Neuron 61, 609&620.

Moldakarimov S., Bazhenov M., and Sejnowski T.J. (2015). Feedback stabilizes propagation of synchronous spiking in cortical neural networks. Proc. Natl. Acad. Sci. (USA).

Murray J.D., Bernacchia A., Freedman D.J., Romo R., Wallis J.D., Cai X., Padoa-Schioppa C., Pasternak T., Seo H., Lee D., and Wang X.J. (2014). A hierarchy of intrinsic timescales across primate cortex. Nat. Neurosci. 17, 1661&1663.

Ogawa T., and Komatsu H. (2010). Differential temporal storage capacity in the baseline activity of neurons in macaque frontal eye field and area V4. J. Neurophysiol. 103, 2433&2445.

Rugh W.J. (1995). Linear System Theory (2nd Edition) (Prentice Hall, New Jersey).

Schmolesky M.T., Wang Y., Hanes D.P., Thompson K.G., Leutgeb S., Schall J.D., and Leventhal A.G. (1998). Signal timing across the macaque visual system. J. Neurophysiol. 79, 3272&3278.

Scholtens L.H., Schmidt R., de Reus M.A., and van den Heuvel M.P. (2014). Linking macroscale graph analytical organization to microscale neuroarchitectonics in the macaque connectome. J. Neurosci. 34, 12192&12205.

Sepulcre J., Liu H., Talukdar T., Martincorena I., Yeo B.T., and Buckner R.L. (2010). The organization of local and distant functional connectivity in the human brain. PLoS Comput. Biol. 6, e1000808.

Shepherd G.M., Stepanyants A., Bureau I., Chklovskii D., and Svoboda K. (2005). Geometric and functional organization of cortical circuits. Nat. Neurosci. 8, 782&790.

Sherrington C.S. (1906). Observations on the scratch-reflex in the spinal dog. J. Physiol. (Lond.) 34, 1&50.

Smith P.L., and Ratcliff R. (2004). Psychology and neurobiology of simple decisions. Trends Neurosci. 27, 161&168.

Sporns O. (2014). Contributions and challenges for network models in cognitive neuroscience. Nat. Neurosci. 17, 652&660.

van den Heuvel M.P., Kahn R.S., Goni J., and Sporns O. (2012). High-cost, high-capacity backbone for global brain communication. Proc. Natl. Acad. Sci. U.S.A. 109, 11372&11377.

Van Essen D.C., Drury H.A., Dickson J., Harwell J., Hanlon D., Anderson C.H., and Van Essen D.C. (2001). An integrated software suite for surface-based analyses of cerebral cortex. J Am Med Inform Assoc 8, 443&459.

Wallisch P., and Movshon J.A. (2008). Structure and function come unglued in the visual cortex. Neuron 60, 195&197.

Wang X.J. (1998). Calcium coding and adaptive temporal computation in cortical pyramidal neurons. J. Neurophysiol. 79, 1549&1566.

Wang X.J. (2001). Synaptic reverberation underlying mnemonic persistent activity. Trends Neurosci. 24, 455&463.

Wang X.J. (2002). Probabilistic decision making by slow reverberation in cortical circuits. Neuron 36, 955&968.

Wang X.J. (2008). Decision making in recurrent neuronal circuits. Neuron 60, 215&234.

Wang X.J. (2013). The prefrontal cortex as a quintessential “cognitive-type” neural circuit: working memory and decision making. In Principles of Frontal Lobe Function, D.T. Stuss, and R.T. Knight, eds. (Cambridge University Press), pp. 226&248.

Wong K.F., and Wang X.J. (2006). A recurrent network mechanism of time integration in perceptual decisions. J. Neurosci. 26, 1314&1328.

## References

Binzegger T., Douglas R.J., and Martin K.A. (2004). A quantitative map of the circuit of cat primary visual cortex. J. Neurosci. 24, 8441&8453.

Cribari-Neto F., and Zeileis A. (2010). Beta regression in R. J. Stat. Softw. 34, 1&24.

Deco G., Jirsa V., McIntosh A.R., Sporns O., and Kotter R. (2009). Key role of coupling, delay, and noise in resting brain fluctuations. Proc. Natl. Acad. Sci. U.S.A. 106, 10302&10307.

Deco G., Ponce-Alvarez A., Mantini D., Romani G.L., Hagmann P., and Corbetta M. (2013). Resting-state functional connectivity emerges from structurally and dynamically shaped slow linear fluctuations. J. Neurosci. 33, 11239&11252.

Gardiner C.W. (1985). Handbook of stochastic methods, vol. 3 (Springer Berlin).

Hilgetag C.C., O’Neill M.A., and Young M.P. (1996). Indeterminate organization of the visual system. Science 271, 776&777.

Markov N.T., Ercsey-Ravasz M., Van Essen D.C., Knoblauch K., Toroczkai Z., and Kennedy H. (2013). Cortical high-density counterstream architectures. Science 342, 1238406.

Markov N.T., Ercsey-Ravasz Gomes M.M., Ribeiro A.R., Lamy C., Magrou L., Vezoli J., Misery P., Falchier A., Quilodran R., Gariel M.A., et al (2014a). A weighted and directed interareal connectivity matrix for macaque cerebral cortex. Cereb. Cortex 24, 17&36.

